# Snapshot of the Atlantic Forest canopy: surveying arboreal mammals in a biodiversity hotspot

**DOI:** 10.1101/2021.03.08.434374

**Authors:** Mariane C. Kaizer, Thiago H.G. Alvim, Claudio L. Novaes, Allan D. Mcdevitt, Robert J. Young

## Abstract

The Atlantic Forest of South America supports some of the greatest terrestrial biodiversity on our planet but is now reduced to only a small extent of its original forest cover. It hosts a large number of endemic mammalian species but our knowledge on arboreal mammal ecology and conservation has been hindered by the challenges of observing arboreal species from ground level. Camera trapping has proven to be an effective tool in terrestrial mammal monitoring, but the technique has rarely been used for arboreal species. Here we generated data on the arboreal mammal community based on canopy camera trapping for the first time in the Atlantic Forest, focusing on Caparaó National Park, Brazil. We placed 24 infrared camera traps in the forest canopy distributed in seven areas within the Park, operating continuously from January 2017 to June 2019. In this time, they accumulated 4,736 camera-days and generated 2,256 sets of pictures and 30 second videos of vertebrates. Canopy camera traps were able to detect arboreal mammals spanning a wide variety of body sizes. The local mammal assemblage comprised of 15 identifiable species, including the critically endangered northern muriqui (*Brachyteles hypoxanthus*) and the buffy-headed marmoset (*Callithrix flaviceps*), and other rare, nocturnal and inconspicuous species. For the first time, we confirmed the occurrence of the thin-spined porcupine (*Chaetomys subspinosus*) in the Park. Species richness varied across sampling areas and forest types. Our findings demonstrate the potential of canopy camera trapping for future surveying efforts to better inform conservation strategies for mammals.

## Introduction

Tropical forest canopies host between half to two-thirds of the terrestrial biodiversity on Earth, yet remain one of the most poorly explored habitats because they are difficult to access (Linsenmair et al., 2001; Lowman, 2009; Lowman et al., 2013). About three quarters of terrestrial forest vertebrates in the tropics, including a great diversity of mammals, are strictly or partially restricted to the arboreal realm (Eisenberg & Thorington, 1973; Kays & Allison, 2001). For many years, tropical arboreal mammals were traditionally inventoried and observed through ground-based methods, which often missed cryptic, fast-moving and nocturnal species (Lowman & Moffett, 1993; Kays & Allison, 2001; Whitworth et al., 2016; Bowler et al., 2017; Moore et al., 2020). These methods are also extremely difficult to implement in remote areas and on a large scale. Recent advances in canopy access techniques (Lowman, 2009) and the incorporation of emerging technologies into conservation efforts (Pimm et al., 2015; Marvin et al., 2016), have been proven to be useful to overcome the aforementioned shortcomings; thereby increasing our knowledge on arboreal mammal species (e.g., arboreal camera trap: Gregory et al., 2014; drones: Kays et al., 2019; passive acoustic recording: Duarte et al., 2018; and environmental DNA (eDNA): Sales et al., 2020).

Identifying effective approaches to assess and monitor the arboreal mammal community is vital to fill the gaps in scientific knowledge and to drive successful management and conservation efforts. Arboreal mammals comprise a great proportion of overall rainforest community biomass, and play fundamental functional roles in the maintenance of the forest ecosystem (Kays & Allison, 2001), including pollination, top-down regulation of prey, folivory, seed dispersal and maintenance of forest carbon storage (Kays & Allison, 2001; Jorge et al., 2013; Bello et al., 2015; Bufalo et al., 2016; Bogoni et al., 2019). Arboreal mammals are more sensitive to habitat disturbance (Whitworth et al., 2019), therefore, anthropogenic impacts may rapidly lead to a loss or decline of wildlife (Dirzo et al., 2014). Consequently, this may cause changes in community composition and functional diversity (Jorge et al., 2013; Dirzo et al., 2014; Bovendorp et al., 2019).

The Atlantic Forest of South America is one of the “hottest” global biodiversity hotspots because it harbours: one of the world’s highest diversity of plants and vertebrates; a high level of endemism; and includes many threatened species (Myers et al., 2000; Laurance, 2009). This originally vast biome (1.5 million km^2^) is now reduced to only ∼12% (∼16,377,472 ha) of its original forest cover, most of which persists as highly fragmented areas smaller than 50-ha (Ribeiro et al., 2009). Over 300 species of mammals occur in the Atlantic Forest biome, with about 30% of the species being endemic (Paglia et al., 2012; Quintela et al., 2020). Although the Atlantic Forest’s mammals are mostly arboreal (Paglia et al., 2012), the majority of studies which have focused on mammals larger than 1kg have been predominantly conducted using ground-based methods, such as transect census and terrestrial camera traps, along with indirect evidence obtained from vocalizations, tracks, faeces and carcasses (Chiarello, 2000; Srbek-Araujo & Chiarello, 2005; Oliveira et al., 2013; Geise et al., 2017). Given that mammal defaunation has been widely documented throughout the Atlantic Forest biome (Canale et al., 2012; Galetti, et al., 2017; Sousa & Srbek-Araujo, 2017; Bogoni et al., 2018, 2020) and will likely continue to increase due to ongoing anthropogenic activities and climate change, there is clearly an urgent need to gather more reliable data on the distribution and population status of arboreal mammals to better inform conservation plans.

Over the last two decades, camera traps have been proven to be an effective non-invasive method to detect rare and elusive species, even in remote areas and over large spatial and temporal scales (Burton et al., 2015; Wearn & Glover-Kapfer, 2019). While camera traps have become a ubiquitous method in ecological studies and conservation programs of terrestrial mammals (Glover-Kapfer et al., 2019), with great potential for global network monitoring (Ahumada et al., 2011; Steenweg et al., 2017). It is only recently that this method has started to be applied to survey arboreal mammals in tropical forest canopies (Olson et al., 2012; Gregory et al., 2014; Whitworth et al., 2016; Bowler et al., 2017; Kaizer, 2019; Hongo et al., 2020; Moore et al., 2020). Here we provide the first study using canopy camera trapping to survey arboreal mammals in the Atlantic Forest. We were particularly interested in the efficiency of this technique to detect a variety of mammals that vary greatly in body size. Furthermore, we wished to assess species richness, community composition and functional traits of arboreal mammal assemblage in the Caparaó National Park, Brazil, particularly given that studies across the globe have found that legally protected areas may harbor higher diversities of mammals (Littlewood et al., 2020). Although the Caparaó National Park is one of the last significant Atlantic Forest remnants (in terms of size) in southeastern Brazil, there is a lack of studies on the vertebrate biodiversity inhabiting the Park and its arboreal mammal community is largely unknown.

## Methods

### Study area

The Caparaó National Park (PNC) is located on the border between the states of Minas Gerais and Espírito Santo, in Southeastern Brazil (20°37’S and 20°19’S and 41°43’W and 41°55’W; Fig. 1). The 31,853 ha of protected area of the park is inserted in the Caparaó massif, which is part of the northern portion of the Mantiqueira mountain range, and stretches for about 40km from north to south, with altitudes ranging between 630m and 2,952m (ICMBIO, 2015). The vegetation types include Mountainous Ombrophilous Dense forest and Mountainous Semideciduous Seasonal forest below 1,500m in altitude, cloud forest between 1,500-1,900m, and high-altitude grasslands above 1900m (Veloso et al., 1991; ICMBIO, 2015). While semi-deciduous Seasonal forest occurs predominantly on the western side of the park, Mountainous Ombrophilous Dense forest is found exclusively in the eastern side (ICMBIO, 2015). The landscape surrounding the PNC is dominated by coffee plantations, pastures and isolated/small forest patches. The climate is humid with a temperate summer according to Koppen’s classification (Alvares et al., 2013). The mean annual temperature is about 19°C in the lower altitudes and 9.4°C in the upper altitudes (Alvares et al., 2013). The mean annual rainfall is around 1,500mm, and the air relative humidity is high (>70%) in almost all months of the year (ICMBIO, 2015). The rainy season occurs from October to April, and a dry and cool season, with monthly rainfall lower than 50mm, from May to September (Alvares et al., 2013; ICMBIO, 2015).

**FIG 1.**
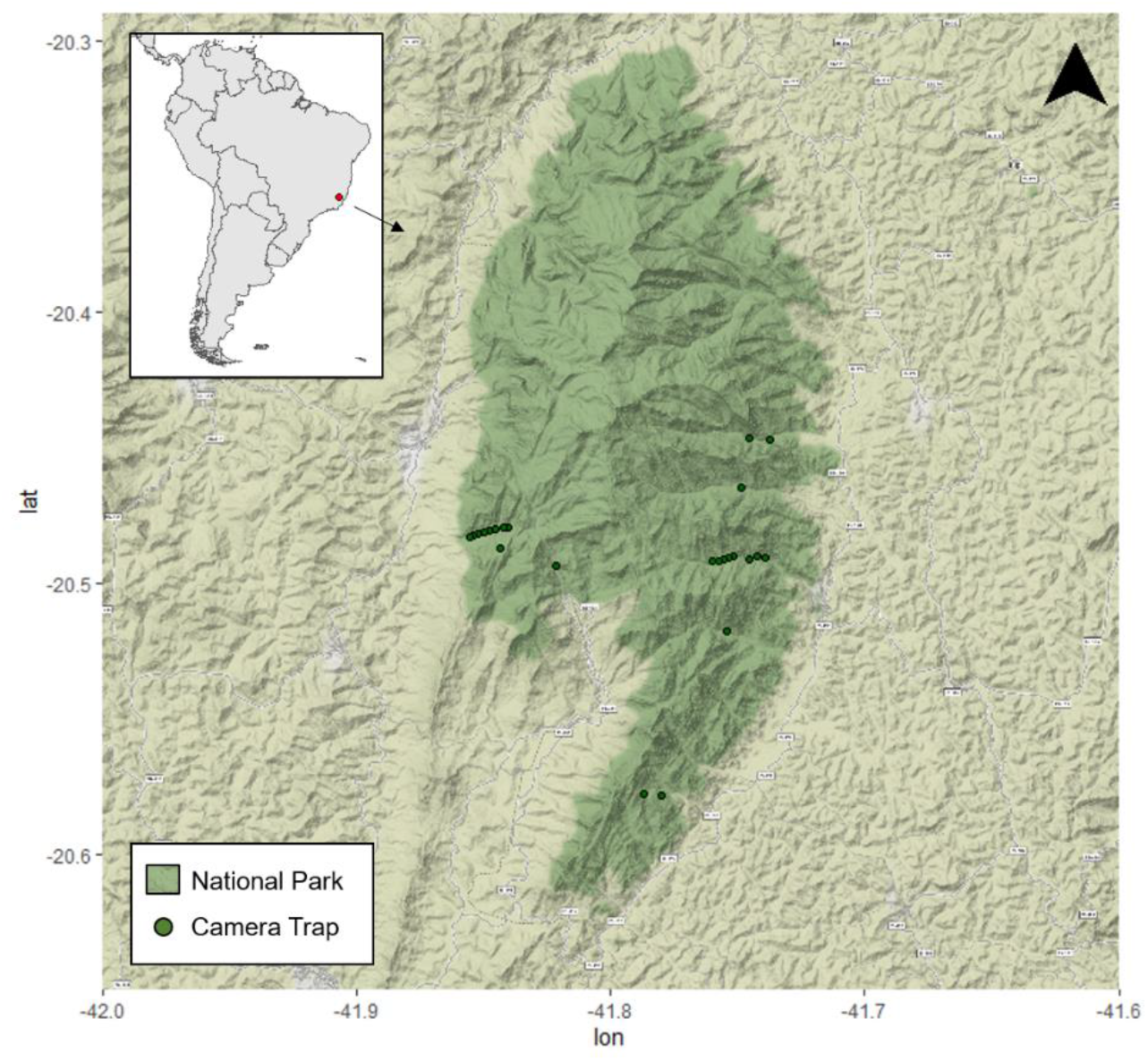
Locations where arboreal camera-trap surveys were conducted in Caparaó National Park, south-eastern Brazil.

### Arboreal Camera Trapping

Infrared camera traps (Bushnell Trophy Cam; Model #119774C) were deployed at PNC from January 2017 through June 2019, as part of a larger project monitoring the resident northern muriqui (*Brachyteles hypoxanthus*) population (Kaizer, 2019). In total, we surveyed 24 sites (each with a single camera trap) in the canopy of Atlantic Forest, distributed in seven distinct valleys within the Park (called: Aleixo, Calçado, Facão de Pedra, Santa Marta, Rio Norte, Rio Preto, and Rio Veado). Survey sites covered an elevation range of 1000–1768m of which 10 sites were on the western side of the park (Montane Semideciduous Seasonal forest) and 14 sites were on the eastern side (Montane Ombrophilous Dense forest). In 2017, eight cameras were placed in each of two valleys (16 cameras in total), one on the west side and one on the east side of the Park, along an elevation gradient (Fig. 1; Kaizer, 2019). In the following years of 2018 and 2019, a further eight camera traps were randomly distributed within the Park (Fig. 1). Arboreal camera traps were placed >250 m apart, strapped to a tree at a mean height of 12.0 m ± 3.1 (range: 7.5-17.0 m) from the ground. Canopy camera trap locations were chosen based on tree connectivity; that is, places where animals could cross the canopy, and considering safe access for climbers (Gregory et al., 2014; Kaizer, 2019; Whitworth et al., 2019). Camera traps were not baited or oriented to east or west (to avoid direct sunlight) and were placed facing along a horizontal branch of the tree or to the branches of an adjacent tree. For a detailed explanation of the arboreal camera trap placement see Chapter 2 in Kaizer (2019). Cameras were active 24h per day and set to hybrid mode (two pictures and one 30 second video), with 10 second intervals between triggers, and low night time LED intensity. Animals were identified on photographs using Wild.ID 0.9.3.1 software (TEAM Network, www.wildlifeinsights.org/team-network). A capture event was defined as a set of two pictures and one 30 second video. To ensure independence between capture events we used a minimum of 1 hour intervals between species-specific records.

Detection rates for arboreal mammals was defined as the ratio of independent events to the number of trap days, which is the number of 24 hour periods from camera placement until the battery ran out or the camera was retrieved, and multiplied by 100 (Rovero & Marshall, 2009). We adopted the camera trap detection rate as a mean to estimate relative abundance of arboreal mammal species (Rovero & Marshall, 2009; Pal et al., 2020). All analyses were run in R v. 3.6.3 environment (R Core Team, 2020).

## Results

Across 24 arboreal camera trap sample points our survey effort totalled 4,736 camera trap days of which 2,151 camera-days were in semideciduous forest and 2,585 camera-days were in ombrophilous forest. Due to camera malfunctioning or battery failure, camera traps were active for a mean of 57.0 days (± 43.6 days SD, range: 1–285). The overall effort resulted in a total of 27,310 trigger events of which 2,256 represented animal species (8.3%), including mammals, birds and lizards.

We obtained 2,200 capture events of arboreal mammals in the canopy of the Atlantic forest, which resulted in 1,396 independent records. From these, we were able to identify 1,216 records (87.2% of the total mammal records), resulting in 15 mammals identified to species-level, two to genus-level and one to family level (Table 1). Unidentifiable records (N=178 records) represented small mammals, including opossums and rodents. The identified arboreal mammals represent 12 families and eight orders (Table 1). Overall, Rodentia was the richest order (N=5 species), followed by Carnivora (N=4) and Didelphidae (N=4), Primates (N=3), Pilosa (N=1) and Chiroptera (N=1).

**TABLE 1.**
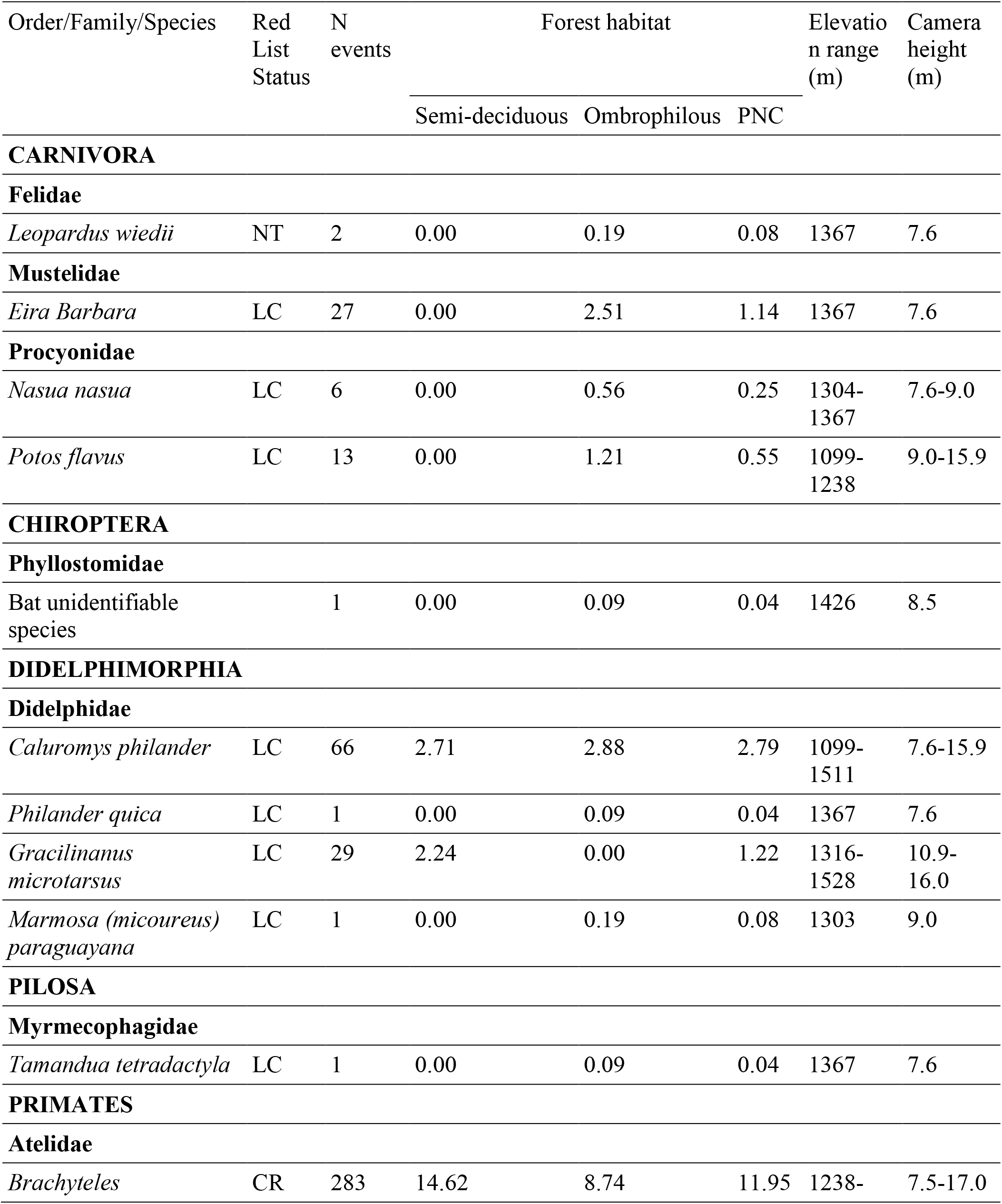
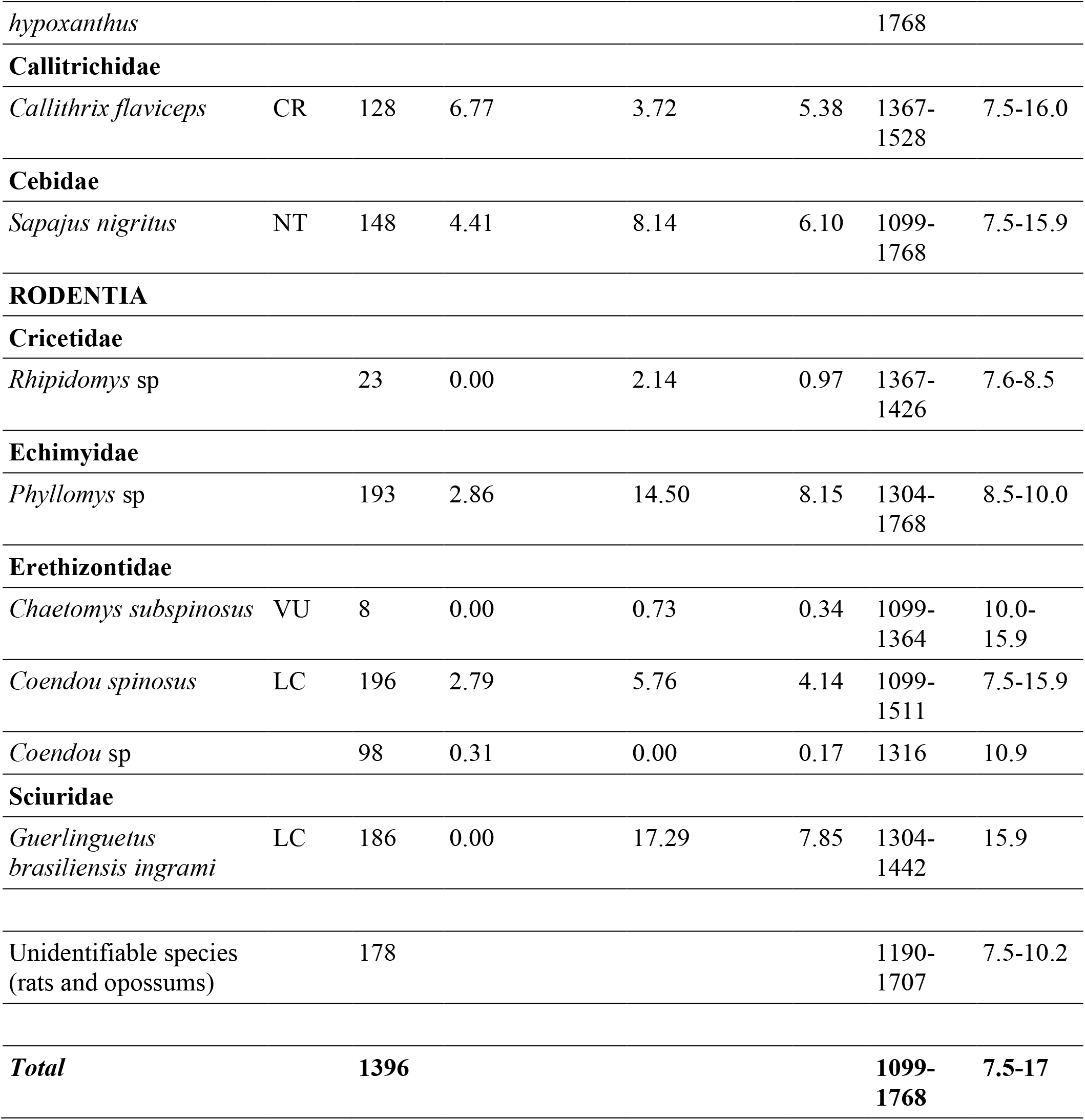
List of mammals recorded by arboreal camera trapping in the Caparaó National Park, Brazil, showing their Red List status (IUCN, 2020), number of independent events, detection rates (independent photographs/100 trap days) in two forest types, elevation range, and camera height range.

Of the mammal species recorded, two are classified as Critically Endangered, one Vulnerable, and two Near Threatened on the IUCN Red List (IUCN, 2020) (Table 1). Seven species, including three primates (*Brachyteles hypoxanthus, Callithrix flaviceps*, and *Sapajus nigritus*), one porcupine (*Chaetomys subspinosus*), one squirrel (*Guerlinguetus b. ingrami*), one tree rat (*Phyllomys* sp), and one opossum (*Gracilinanus microtarsus*) are endemic to the Atlantic Forest. The detection of the thin-spined porcupine (*C. subspinosus*) was the first confirmation of the occurrence of the species in the park and in the western portion of the Espirito Santo State (Giné & Faria, 2018). Arboreal camera traps detected mammals spanning a wide range of body sizes (Table 1, Plate 1), nine of them heavier than 1kg. The largest bodied species detected was the primate, the northern muriqui (*B. hypoxanthus*, 13 kg), and the smallest-bodied mammals were *G. microtarsus* (12-52 g), *Rhipidomys* sp (50-90g), *Marmosa m. paraguayana* (120-175 g), *G. b. ingrami* (125-216 g), *Caluromys philander* (140-390 g), *P. pattoni* (212 g), *Philander quica* (220-680 g) and *C. flaviceps* (400 g) (Paglia et al., 2012; Faria et al., 2019). The majority of mammal species were arboreal, but five mammals were scansorial (*G. b. ingrami, M. m. paragiayana, P. quica, Tamandua tetradactyla* and *Leopardus wiedii*) and two were terrestrial (*Eira barbara* and *Nasua nasua*).

**PLATE 1.**
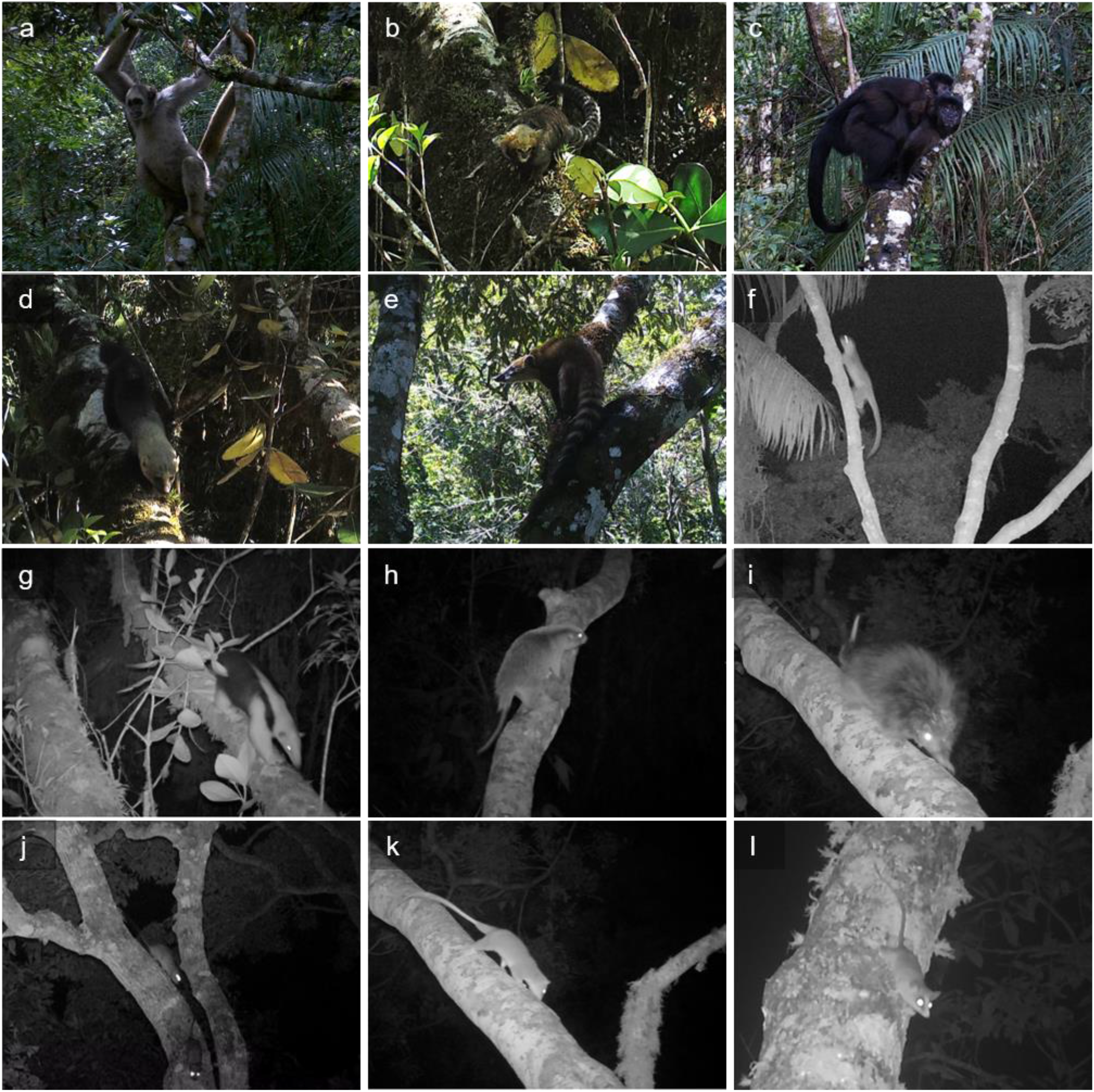
Some arboreal mammals photographed by camera trap in the canopy of Atlantic Forest, Caparaó National Park, Brazil: a) northern muriqui (*Brachyteles hypoxanthus*), b) buffy-headed marmoset (*Callithrix flaviceps*), c) black-horned capuchin (*Sapajus nigritus*), d) tayra (*Eira barbara*), e) South American coati (*Nasua nasua*), f) kinkajou (*Potos flavus*), g) southern tamandua (*Tamandua tetradactyla*), h) thin-spined rat (*Chaetomys subspinosus*), i) spiny tree porcupine (*Coendou spinosus*), j) rusty-sided Atlantic tree-rat (*Phyllomys* sp), k) bare-tailed woolly opossum (*Caluromys philander*), l) Brazilian gracile opossum (*Gracilinanus microtarsus*).

Species richness and mammal community composition also varied between forest types. Arboreal cameras documented greater species richness and functional diversity in the community in ombrophilous dense forest (Table 1, Fig. 2). Furthermore, 10 species were recorded exclusively in the montane ombrophilous dense forest, and only one species was exclusively recorded in the semi-deciduous forest (Table 1, Fig. 2). Regarding the functional traits of the community, species richness of frugivore-omnivore species and frugivore-folivore was higher in both forest types. However, two functional groups were missing in the semi-deciduous forest (e.g. carnivore and myrmecophagy).

**FIG 2.**
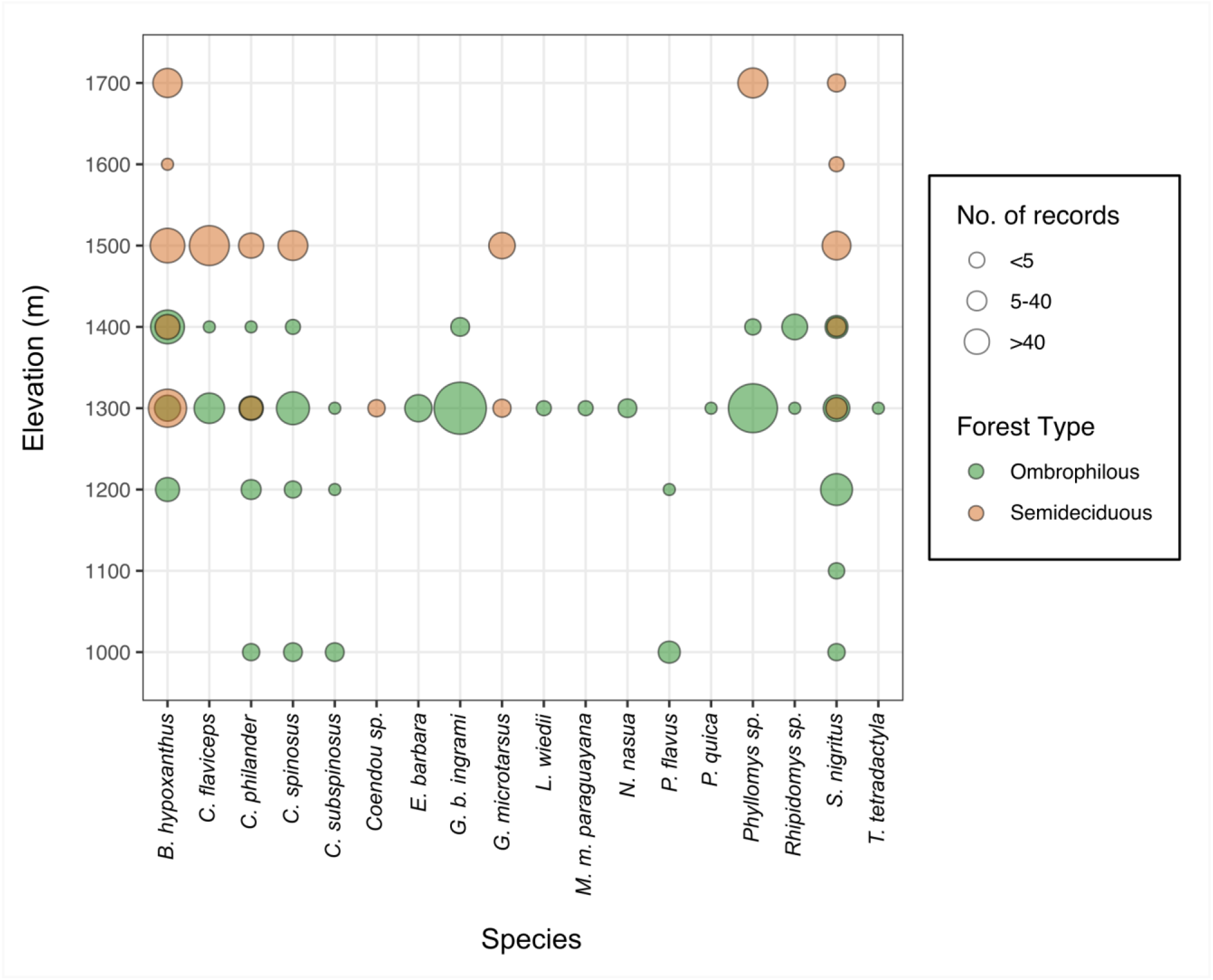
Bubble graph representing presence-absence and categorical values of the number of records in each forest type (Semi-deciduous forest and Ombrophilous Dense forest) for arboreal camera traps for each species identified across an elevational gradient in Caparaó National Park, Atlantic Forest, south-eastern Brazil.

Overall detection rate estimates for the mammals detected by arboreal camera traps were highly variable ranging from 0.04 for *M. m. paraguayana, Philander quica, T. tetradactyla* and Phyllostomidae bat to 11.95 for *B. hypoxanthus* (Table 1). Between forest types, detection rate estimates for mammals in semi-deciduous forest ranged from 2.24 for *G. microtarsus* to 14.62 for *B. hypoxanthus*, while in ombrophilous forest detection rates ranged from 0.09 for *M. m. paraguayana, P. quica, T. tetradactyla* and Phyllostomidae bat to 17.29 for *G. b. ingrami*. The northern muriqui (*B. hypoxanthus*) and the black-horned capuchin (*S. nigritus*) were detected widely (20 and 17 canopy sampling locations, respectively) and across an elevation gradient (Fig. 2). Whereas, the *M. m. paraguayana, P. quica*, Phyllostomidae bat and *T. tetradactyla* were detected only once, and *L. wiedii* twice during the survey period. The kinkajou (*P. flavus*) and the thin-spined porcupine (*C. spinosus*) were detected at two and three sampling locations close to streams in ombrophilous forest type, in elevation of up to 1238 m and 1364 m, respectively (Fig. 2). Regarding the functional traits, overall mean relative abundance was highest for folivore (8.1, N=1), frugivore-gramivore (7.9, N=1), frugivore-folivore (5.5 ± 5.9 SD, N=3) and frugivore-insectivre-gramivore (5.4, N=1) species. Between forest types, frugivore-insectivore-gomivore (6.8, N=1), frugivore-folivore (5.9 ± 7.71 SD, N=3), and folivore (2.9, N=1) species presented the highest mean relative abundance in semideciduous forest. Whereas, frugivore-gramivore (17.3, N=1) folivore (14.5, N=1), and frugivore-folivore (5.1 ± 4.0 SD, N=3) species were most abundant in ombrophilous dense forest.

## Discussion

We examined species richness, community composition and functional traits of arboreal mammals in the Atlantic Forest canopy. As far as we are aware, this is the first study using arboreal camera trapping to assess mammal assemblage in the canopy of this biodiversity hotspot (but for low camera trap height studies see Kierulff et al. (2004) and Oliveira-Santos et al. (2008)). Our results demonstrate the efficiency of this method to detect arboreal mammals of distinct body sizes, and to detect rare and highly cryptic species such as the buffy-headed marmoset, tree-rat and thin-spined porcupine. Furthermore, we compiled evidence that the protected area maintains an important species richness and functional community of arboreal mammals, including the largest arboreal seed disperser (*B. hypoxanthus*). Results also indicate a greater diversity of mammals in the ombrophilous forest type. Nonetheless, the real species richness of arboreal mammal assemblage in each forest type, and thus in our entire study site, may be underestimated by canopy cameras traps alone due to the ecology of some species. For example, terrestrial or scansorial species, such as *Eira barbara, M. m. paraguayana, Nasua nasua, T. tetradactyla*, and *L. wiedii*, were detected only at ombrophilous forest but their presence or absence may be under-documented when only using arboreal camera traps.

The number of arboreal mammals documented in this study is comparable to species richness reported in different arboreal camera trapping studies conducted in other tropical rainforest sites. In the Amazon forest of Peru, species richness of arboreal mammals detected in canopy camera traps ranged from 18 to 24 species (18 species: Whitworth et al., 2016; Bowler et al., 2017; 20 species: Gregory et al., 2014; and 24 species: Whitworth et al., 2019). In the west African rainforest, arboreal cameras recorded 18 arboreal mammals at species level and 1 at genus level in Boumba-Bek and Nki National Parks, Cameroon (Hongo et al., 2020), and 15 arboreal mammals at species level and 2 at genus level in Nyungwe National Park, Rwanda (Moore et al., 2020). Major differences to these studies, in relation to our study, relate to the high species richness of primates found in those habitats, and also squirrel species in Rwanda. At least six species of primates are known to occur in the rainforest of Caparaó National Park (Culot et al., 2019), with three of them detected in this study. The absence or non-detection of the other three species (*Callicebus nigrifrons, C. personatus* and *Alouatta guariba*) may be related to a recent yellow fever outbreak, which caused the death of more than 5,000 non-human primates in the Atlantic Forest (Bicca-Marques et al., 2017). The last sighting of *A. guariba* and *C. nigrifrons* in our study area was reported by the lead author (MCK) between December 2016 and March 2017, respectively. This coincided with the period of the most severe yellow fever outbreak in southeastern Brazil (Faria et al., 2018). Our results suggest that further research is necessary to evaluate the current population status of these primate species in the park to fully evaluate the impact of yellow fever.

Defaunation and collapse of functional diversity of the mammal community have been reported throughout of Atlantic Forest biome (Jorge et al., 2013; Galetti et al., 2015; Galetti et al., 2017; Bogoni et al., 2018, 2020) and this has occurred even in large protected areas (Canale et al., 2012). Coupled with the historical habitat loss and fragmentation of the Atlantic Forest, historical and recurrent hunting pressure are the major drivers of mammal defaunation and community composition changes (Jorge et al., 2013; Bogoni et al., 2018). Based on studies from 1983 to 2015, Bogoni et al., (2017) reported a mean species richness of 14.7 for mammal assemblages in Atlantic Forest fragments, when considering only species >1 kg. Our results reported the occurrence of nine species larger than 1 kg in the forest canopy in our study. The majority of these species are frugivore-omnivore and frugivore-folivore, which play an important role as seed dispersers in tropical forests and in nutrient cycling. The northern muriqui, for example, exert a pivotal role in the dispersal and recruitment of large-seed species, which has consequences for key ecosystem services such as carbon stock (Bufalo et al., 2016). Frugivore-folivore species also have a positive correlation with dung-beetles species richness across the Atlantic Forest, playing a fundamental role in nutrient cycling, quality of soil and detritivore food webs (Nichols et al., 2008; Bogoni et al., 2019). Regarding the small body-sized species (<1kg), our study revealed the occurrence of key species, such as large rodents and marsupials from the genera *Phyllomys* and *Caluromys*, which are usually the first groups to disappear from disturbed habitats (Chiarello, 1999).

Our results reinforce the important role of protected areas for mammal conservation (Littlewood et al., 2020). It is estimated that less than 3% of Atlantic Forest remnants are suitable habitat to host the thin-spined porcupine (*C. subspinosus*; Bonvicino et al., 2018). The occurrence of this species in the Park is therefore pivotal for its persistence in the long-term. The first documentation of the tree-rat (*Phyllomys* sp.) also demonstrates the potential of the PNC to host rare species. Although we were not able to confirm this record to species-level, Faria et al. (2016) recently recorded the rare *Phyllomys lundi* in a private reserve ca. 20 km from our study sites. This endangered species is known from only three locations in the entire Atlantic Forest biome, so confirmation of its presence would expand its range to a new location (Bonvicino, 2018; Faria et al., 2016). The Caparaó National Park has also been indicated to be one of the four priority areas for conservation of the critically endangered northern muriqui (Melo et al., 2018). This distinct population is also of high importance because it inhabits the largest altitudinal range of the species (up to 2,000 m above sea level; Strier et al., 2017). By using arboreal camera trapping, we were able to provide valuable information on this species across an elevational gradient, including in high elevations and topographic slopes, where accessibility to perform surveys by traditional ground-based methods is limited. Furthermore, the northern muriqui was only recently discovered to occur on the west side of the park (Kaizer et al., 2016), and its occurrence in ombrophilous forest was previously only reported in a few locations (Mendes et al., 2005). Our findings provide new records for the occurrence of this species in two new sites within the park (valleys of Rio Preto and Rio Norte). This demonstrates the importance of this protected area to safeguard this evolutionarily distinct and globally endangered species (EDGE species; Isaac et al., 2007). Despite this, the high detection rate of northern muriqui in our study site seems to reflect the large home range of the species (Dias & Strier, 2003; Lima et al., 2019), which was detected along an array of arboreal camera traps in the same valley.

Although ∼12% of the records of mammals in this study represented small mammals that were not identifiable, including bats, rodents and opossums, our results demonstrate the ability of arboreal cameras to detect smaller bodied species. The record of *Phyllomys* sp. is an example of the potential of arboreal cameras to detect elusive and arboreal species that are usually difficult to collect using small mammal traps (Faria et al., 2016; Bonvicino et al., 2018). However, since most of these records were nocturnal, thus hampering the recognition of some species, the use of camera traps with white flash functionality might increase the potential and effectiveness of this method (Bowler et al., 2017). The configuration of the cameras to hybrid mode (i.e. to record a short video following a still picture) also increases the likelihood of distinguishing species. Furthermore, this provides better documentation of fast-moving species, such as squirrels and marmosets (also indicating the number of individuals and the potential for collecting data on species behavior; Caravaggi et al., 2020).

To date, most of the studies reporting arboreal mammal assemblage in remnants of the Atlantic Forest have been based on ground-based surveys. Our findings demonstrate the potential of arboreal camera trapping to register rare, nocturnal and cryptic species that are prone to false negatives obtained by ground-based methods (Olson et al., 2012; Whitworth et al., 2016; Bowler et al., 2017; Moore et al., 2020). Considering the habits of some scansorial and terrestrial species, we suggest that arboreal camera traps should be paired with terrestrial cameras, which would reduce the likelihood of failing to detect these species. This would therefore provide a better snapshot of the entire mammal assemblage. In addition, our results illustrate the role of the Caparaó National Park as a stronghold for the conservation of rare and threatened mammalian species endemic in one of the most important global biodiversity hotspots. We encourage future studies over larger spatial and temporal scales, which aim to explore trends in species composition and functional diversity of the entire mammalian community and in conjunction with emerging biomonitoring technologies (e.g. eDNA; Sales et al., 2020). This will provide a more complete understanding of how functional diversity and ecosystem multifunctionality is conserved, and to better inform evidence-based conservation strategies for this protected area.

## Author contributions

Study design and fieldwork: MCK, CLN and THGA; camera trap data processing: CLN and MCK; data analysis and writing the article: MCK, ADM and RJY.

## Acknowledgements

The authors thank the Brazilian Ministry of Environment/SISBIO for authorizing research in the Caparaó National Park and park managers for logistical support. We are grateful to Francisco Homem, Leandro Moreira, Rodrigo Silva and Viviane Sodré for fieldwork assistance, Aryanne Clyvia and Daniel Ferraz for logistical support, and Guilherme Garbino, Michel Faria and Rayque Lanes for the identification of small mammals. We acknowledge the Brazilian Ministry of Education/CAPES (BEX 1 298/2015-01) for the award of a PhD studentship to MCK, Idea Wild and Conquista Montanhismo for equipment grants, and Conservation Leadership Programme (N. 12455) and MBZ Conservation fund (N. 162512917) for funding support to the Caparaó Muriqui Project, which this work is part of. Finally, thanks to the Conservation Leadership Programme for supporting MCK to be part of the Writing for Conservation Workshop, and the National Geographic Society for supporting MCK as an Early Career National Geographic Explorer.

## Conflicts of interest

‘*None*’

## Ethical standards

This research was approved by the Caparaó National Park (ICMBio/SISBIO No: 49062) and University of Salford (STR1718-14) and abided by the *Oryx* guidelines on ethical standards.

## Notes

### Competing Interest Statement

The authors have declared no competing interest.

